# PMAT enhances sexual dimorphism of fear behaviors and facilitates female mice’s generalized contextual fear extinction

**DOI:** 10.1101/2025.08.19.671133

**Authors:** Aliyah J. Ross, Lauren R. Scrimshaw, Lauren N. Stoner, Jasmin N. Beaver, Grace A. Sonick, Lee M. Gilman

## Abstract

Enhanced signaling of dopamine and/or serotonin during highly arousing situations can be reduced or terminated by monoamine transporters. One such transporter, plasma membrane monoamine transporter (PMAT, *Slc29a4*), attenuates both dopamine and serotonin signaling. An absence of selective pharmacological inhibitors means genetically modified mice constitutively deficient in PMAT remain the best tool with which to study PMAT’s organism-level functional effects. Fear conditioning is a high arousal process. Generalization of fear is an evolutionarily advantageous process, whereby information learned from one experience is applied to other new but similar encounters. Pathological fear generalization, in contrast, is a core feature of most anxiety disorders. Given our previous findings indicating PMAT function reduces male mice’s context fear and enhances extinction of female mice’s cued fear, we hypothesized that PMAT would similarly reduce generalization (i.e., enhance discrimination) of context and cued fear in male and female mice, respectively. Our context and cued fear conditioning experiments in adult PMAT wildtype (+/+) and heterozygous (+/−) male and female mice partially supported our hypotheses. We discovered PMAT functions to facilitate extinction of contextually generalized fear, plus subsequent extinction of context-specific fear, selectively in females. Moreover, when specific fear cues or contexts were temporally presented before cues or contexts that were similar enough to make generalization possible, PMAT enhanced biological sex differences. Growing evidence reports common PMAT polymorphisms elicit measurable effects when PMAT function is reduced. Thus, we suspect future experiments may reveal positive associations between PMAT polymorphisms and risk for anxiety disorder symptoms, particularly in people assigned female at birth. Inclusion of these genetic variations in pharmacogenomic analyses may prove therapeutically beneficial.

## Introduction

During stressful or otherwise highly arousing (e.g., memorable) events, neuronal release of monoamines like dopamine and serotonin, is elevated^1–6^. Brain dopamine and serotonin signaling, in part, is regulated by uptake transporters^7–9^ [see reviews^10,11^]. Such monoamine uptake transporters are typically classified into one of two categories: high affinity, low capacity (i.e., uptake 1); and low affinity, high capacity (i.e., uptake 2). Uptake 1 monoamine transporters have relatively high selectivity for their preferred substrate(s), with affinities (*K*_m_) ranging between 0.08–5.2 μM^12–20^. Uptake 1 have a lower transport capacity, with rates of 0.006-0.300 nmol/min/mg protein for dopamine transporter (DAT, *Slc6a3*)^13,14^ and 0.001-0.012 nmol/min/mg protein for serotonin transporter (SERT, *Slc6a4*)^12,13^. In contrast, the affinity constants for uptake 2 monoamine transporters span from 80-2100 μM for dopamine and serotonin^21–26^, and transport rates vary from 0.42-18.2 nmol/min/mg protein^21–24,27^. Given the abundance of drugs, both legal and illicit, that act upon the uptake 1 transporters DAT, SERT, and norepinephrine transporter (NET, *Slc6a2*), far more research has historically focused upon uptake 1 as opposed to uptake 2 transporters. Indeed, drugs selective for uptake 2 transporters remain elusive^28–30^ (see reviews^31,32^), hindering studies and thereby limiting understanding of their functional contributions. Nevertheless, increasing evidence collected despite a lack of uptake 2-selective pharmacological tools indicates that these transporters are influential in a range of experiences, from olfaction^33^; to drug absorption, tolerance, and response^34–40^; to emotion-related processes and diseases^30,41–43^. Considering the higher selectivity and lower transport rates of uptake 1, the contribution of uptake 2 during elevated signaling circumstances makes physiological sense.

Dopamine and serotonin signaling is enhanced during high arousal situations, and these monoamines contribute to formation and expression of memories^1–5,44^ (see reviews^45,46^). It follows that changes in uptake 2-mediated transport of dopamine and serotonin likely influences learning and memory of high arousal circumstances. Plasma membrane monoamine transporter (PMAT, *Slc29a4*) is one such uptake 2 transporter. PMAT preferentially transports dopamine (K_m_=201-466 mM) and serotonin (K_m_=82.9-283 mM)^21,23,27^ over other monoamines like norepinephrine (K_m_=1078-2606 mM)^21,27^. Further, comparing PMAT transport rates (0.750-22.4 nmol/min/mg protein for DA, 1.26-14.2 nmol/min/mg protein for 5-HT)^21,23,27^ to those published for DAT^13,14^ or SERT^12,13^, respectively, indicates that at minimum PMAT transports dopamine and serotonin 2.5 to 105 times faster (possibly up to 3700-14,200 times faster).

Because selective PMAT inhibitors are not presently available, little remains understood regarding how PMAT function contributes to humans’ neurophysiology or behavior. Several rare *SLC29A4* polymorphisms have been identified, but recent studies of more common *SLC29A4* polymorphisms indicate reductions in functional PMAT likely exert detectable effects in people^30,37,38,40,47,48^. For example, *SLC29A4* single nucleotide polymorphisms rs4724512, rs3889348, rs149798710, rs151039853, and rs17854505 are implicated in impaired intestinal absorption of metformin to an extent that drug adherence, response to metformin, and type II diabetes symptoms are all adversely affected^37,38,40,47,48^. Considering the affinity of PMAT for metformin (K_m_=1,320 mM)^49^ is markedly lower than its affinity for dopamine or serotonin, it is plausible that *SLC29A4* polymorphisms result in meaningful brain monoamine signaling changes, particularly when release is elevated. For example, a recent phenome-wide association analysis flagged *SLC29A4* as a key “druggable gene” for mitigating postpartum depression risk^30^.

Considering no commercially available drugs selectively inhibit or otherwise target PMAT, we and others have capitalized upon the Wang lab’s generation of genetically modified mice with constitutively reduced or ablated PMAT function^50^. Using these mice, we have found that PMAT function modestly enhances anxiety-related behaviors across sexes, sex-selectively enhances (males) or reduces (females) active coping behaviors depending upon stressor type, and attenuates context fear expression in males subsequent to preceding stressful experiences^41–43^. PMAT function may also mildly facilitate the time course of cued fear extinction in females, specifically during initial extinction learning^42^. These sex- and stress-specific effects of PMAT function during fear, coping, and anxiety-related behavior tests in mice, combined with evidence that *SLC29A4* polymorphisms influence people’s health, suggest PMAT could be affecting symptoms and/or risk of anxiety disorders.

Fear generalization is a cross-species phenomenon where learned fear from one stimulus is transferred to a novel stimulus. This natural behavior can reduce cognitive load by enabling one learning experience to be applied to multiple future situations. For instance, if someone were to get stung by a hornet, they would subsequently try to avoid other flying insects (e.g., houseflies, dragonflies, etc.) to protect themselves. Sometimes, however, fear generalization becomes too broad, or otherwise reaches a pathological state. *Pathological* fear generalization is a key symptom of anxiety disorders^51–54^. For example, if someone is bit by a pug as a child, as they age they may develop a fear of dogs of all breeds, from chihuahuas to golden retrievers, even though these look quite distinct from the pug that bit them. Indeed, anxiety disorders like specific phobias, social anxiety disorder, and generalized anxiety disorder are characterized by hallmark symptoms of pathological fear generalization^55–60^. The World Health Organization reported that, in 2019, over 300 million people globally were suffering from anxiety disorders^61^, making these some of the most prevalent mental health conditions worldwide.

Fear generalization is one end of a spectrum; at the other end is fear discrimination. Fear discrimination occurs when a distinction between two similar stimuli is learned, such as distinguishing a fire alarm from a tornado alarm^62^. These two intertwined behaviors have been studied in humans to evaluate their relationships to each other and to anxiety disorders^55–59^. Fear generalization to a perceived threat is consistently increased and more prominent in those with anxiety disorders than those without^52,55–57^. Additionally, fear generalization is expressed differently depending on the type of anxiety disorder^56–58,60^, and symptoms can be exacerbated by particular cues and/or environments. Accordingly, experiments in rodents have evaluated putative neurobiological mechanisms influencing fear generalization and fear discrimination in rodents through both context^41,63–65^ and cued^64,66,67^ paradigms. Contextual paradigms evaluate how a learning (training) environment can be distinguished from, or generalized with, a novel environment. This enables assessment of broader context awareness and encoding, and can inform us how situations like dentists’ exam rooms generate (generalization) or alleviate (discrimination) anxiety symptoms when distinctive in location, arrangement, décor, music, etc. Meanwhile, cued paradigms use two distinct cues – often auditory tones – with only one ever coinciding with an aversive stimulus. Such an approach is directed towards understanding aversive associations formed with discrete cues that are often independent of context. For instance, the voice of a bully could terrorize a person whether they hear it in a classroom, grocery store, community center, or workplace; but the voice of strangers may (generalization) or may not (discrimination) elicit a similar psychophysiological reaction. This study utilized both context and cued paradigms to evaluate how PMAT’s function, and reductions thereof, affects fear generalization and discrimination processes using mice.

Understanding PMAT’s role in fear generalization and fear discrimination processes will give us better understanding of the etiology of anxiety disorders and symptoms that accompany them. Due to PMAT’s role in male mice’s fear expression and stress coping behaviors, we hypothesized PMAT function constrains fear generalization and facilitates fear discrimination when males recall aversive contextual experiences^41,42^. Conversely, we hypothesized that in females, PMAT’s function would be more apparent through improved fear discrimination of auditory cues^42^.

## Materials and Methods

### Animals

All experiments used adult (≥90 days old) male and female PMAT-deficient mice on a C57BL/6J background bred in-house. These mice were developed at the University of Washington by Dr. Joanne Wang^50^ and are bred and used under a material transfer agreement between Kent State University and the University of Washington. Mice were group housed (2-5 per cage) within the same sex in cages containing 7090 Teklad Sani-chip bedding (Envigo, East Millstone, NJ, USA). All mice had *ad libitum* access to water and LabDiet 5001 rodent laboratory chow (LabDiet, Brentwood, MO, USA). Mouse housing rooms were maintained on a 12:12 light:dark cycle, with lights at 07:00 local standard time. Room temperature was maintained at 22 ± 1°C and 40 ± 10% relative humidity. Male and female mice were run in experiments separately, with males always run before females. This was done to minimize males’ behavior from being confounded by the presence of females’ pheromones^68–70^. Kent State’s Institutional Animal Care and Use committee approved all experiments, and conditions that adhered to the National Research Council’s Guide for the Care and Use of Laboratory Animals, 8^th^ Ed.^71^. Power analyses for averaged context and cued comparisons (each involving 2 measurements) were performed *a priori* (repeated measures, within-between interactions; f=0.025; α=0.05; 1−β=0.80; corr=0.5; ε=1; G*Power v 3.1.9.6^72^), and indicated n=12 per sex and genotype (i.e., 4 groups).

### Context fear conditioning

Mice underwent context fear conditioning^42,63,73,74^ in Actimetrics chambers (internally, 18.7cm W x 20.6cm D x 20.1cm H; externally, 21.3cm W x 22.9 cm D x 25.7 cm H; Lafayette, IN) composed of two clear acrylic walls opposite each other, and joined by two aluminum walls. The training context (Context A), used both for context fear training and for testing context fear expression, consisted of a black and white striped background, shock grid floor, visible house light, and specific cleaning solution scent cue (70% ethanol). The novel context (Context B), used for testing context fear generalization, consisted of a black background, smooth floor, infrared light, and a different cleaning solution scent cue (Windex®). Freezing, i.e., absence of all movement except breathing, was quantified in real time using FreezeFrame 6 software (Actimetrics, Lafayette, IN). FreezeFrame 6 was also used to administer shocks during context training.

#### Context fear training

All mice underwent context fear training on day 0. Training involved five, 1s, pseudo-randomly administered scrambled foot shocks at 0.4 mA. Training lasted 6 min, with foot shocks administered at 137, 186, 229, 285, and 324 s. Data were exported in 5 s bins; post-shock freezing was calculated as average percent freezing over 30 s (i.e., six, 5 s bins), starting immediately after the 5 s bin during which each foot shock occurred.

#### Context fear expression and generalization testing

Mice were tested 4 wks after training (day 28) then again 48 h thereafter (day 30). Context fear expression was tested in the training context (A), whereas context fear generalization was tested in the novel context (B). Testing in these two contexts was counterbalanced, meaning mice were either tested for fear expression at day 28, followed on day 30 by testing for fear generalization (A➜B); or mice were tested for fear generalization at day 28, followed on day 30 by fear expression testing (B➜A). Within sex and genotype, assignment to testing context order was systematically randomized to ensure acquisition confounds were unlikely to affect testing outcomes. Context testing lasted 10 min each day, and no shocks were administered on any testing day. Freezing was recorded in 30 s bins for the duration of each 10 min test. These testing data were analyzed both as a time course across all 10 min, and as the average percent freezing during minutes two through six across those respective ten, 30s bins^42,63,73^.

### Cued fear conditioning

Cued fear conditioning^42,66^ was performed in the same Actimetrics chambers as for context fear conditioning. Cued fear training likewise used the same Context A as described for context fear conditioning. Testing of cued fear occurred in a novel context, similar to that described for context fear conditioning, save that floors were stainless steel with circular holes. Computer audio output volume was always set at 50% in Windows 11 Home (Microsoft; Redmond, WA). FreezeFrame 6 software was used to administer foot shocks and all auditory tones, as well as to measure freezing behavior. Two auditory cues were utilized: one tone at 7.5 kHz (paired conditioned stimulus, CS+), and one tone at 2.0 kHz (unpaired conditioned stimulus, CS−). Auditory cues were always played for 30s, and CS+ and CS− tones never coincided. The amplitude within FreezeFrame 6 for CS+ tones was set at 80, while for CS− tones it was set at 14. Before training occurred, decibels in each chamber were measured and ranged between 66-74dB for both tones to ensure comparable tone amplitude.

#### Cued fear training

All mice underwent training on day 0 in the training context. During training, mice were pseudo-randomly presented with five CS+ and five CS− tones. CS+ tones were played at 130, 300, 500, 710, and 910s, and each of these were paired with a 1s co-terminating shock at 0.4 mA. CS− tones were played during 210, 410, 650, 820, and 1000s, and were never paired with any aversive stimulus. Freezing was analyzed as the percent freezing during the 30s tones presented for each CS+ and CS−.

#### Cued fear expression testing, extinction training, and extinction retention testing

Mice experienced expression testing and extinction training on day 2, followed by extinction retention testing on day 4. No shocks were administered during any cued fear testing. Testing occurred in the same novel context as described for context fear testing, except the floor was stainless steel with circular holes with 0.635cm diameters. During day 2 (cued fear expression testing and extinction training), a total of 30 CS+ and 5 CS− tones were played over 37 min. CS+ tones were played during 120, 180, 240, 300, 360, 480, 540, 600, 660, 720, 840, 900, 960, 1020, 1080, 1200, 1260, 1320, 1380, 1440, 1560, 1620, 1680, 1740, 1800, 1920, 1980, 2040, 2100, and 2160s. CS− tones were played at 420, 780, 1140, 1500, and 1860s^66^. For day 4 (extinction retention testing), 15 CS+ and 5 CS− tones were presented across 22 min. CS+ tones were played at 120, 180, 240, 300, 360, 480, 540, 600, 660, 720, 840, 900, 960, 1020, and 1080s. Similarly, CS− tones were played at 420, 780, 1140, 1500, and 1860s. Freezing during each day’s five CS− tones was recorded and graphed individually, or averaged to create a single CS− freezing value each for days 2 and 4. Freezing during the 30 (day 2) or 15 (day 4) CS+ tones presented were averaged in sequential groups of three to evaluate time course of freezing across 10 (day 2) or 5 (day 4) points. For comparison with the single averaged CS− freezing value per mouse, described above, freezing for the first five CS+ tones were averaged per mouse^66^.

### Genotyping

Mice were weaned at postnatal day 21, at which time ear punches were taken for identification and genotyping. Genomic DNA was extracted from ear punches by digesting the 2 mm punches with Proteinase K (Roche, Basel, Switzerland) dissolved to 0.077% w/v the day of extraction with STE buffer (100mM Tris, 0.2% SDS, 200mM NaCl, and 5mM EDTA, pH=8.5)^41,43,50^. After a 2 h incubation at 55°C, DNA from the resulting supernate was precipitated with isopropanol, then pelleted DNA was reconstituted in TE buffer (10 mM Tris, 0.1 mM EDTA, pH=8). DNA amplification reactions occurred in 1X PCR buffer containing 1.74mM MgCl_2_ and 34.7 mM dNTPs, 0.39 mM of each primer, and 0.20 mL of Platinum Taq (Invitrogen, Carlsbad, Ca, USA) per 25.6 mL reaction. Sequences of primers, designed by Duan and Wang^50^, have been published previously^41,43,50^. Cycling conditions: 95 °C for 5 min; 34 cycles of 94 °C for 30 s, 59 °C for 30 s, and 72 °C for 90 s; 72 °C for 5 min; hold at 4 °C. Products were visualized using a 1% agarose gel electrophoresis ran at 150 V for 30 min. DNA amplicon sizes were referenced using a 1 kb DNA ladder (Invitrogen), with knockout alleles at 447 bp and wildtype alleles at 847 bp^41,43,50^. All genotypes were re-verified following experiment completion.

### Statistical analyses

Analyses were performed with IBM SPSS Statistics (v 29.0.1.1 (244), IBM, Armonk, NY, USA). The threshold for significance was set *a priori* at p<0.05. Trends (p<0.10) that did not reach this threshold were only acknowledged in text if the associated partial η^2^ (η_p_^2^) was ≥0.060. Cued data were analyzed over time for each conditioning stage: 1) training; 2) fear expression testing and extinction training; 3) extinction retention testing. Context data were likewise analyzed over time at each stage: 1) training; 2) first testing; 3) second testing. These time course analyses were performed using two-way repeated measures (RM) general linear models (GLMs; time × sex × genotype), with Greenhouse-Geisser corrections for within-subjects analyses. Stage 2 cued averages of the first five CS+ and five CS− tones^66^ were also analyzed with two-way RM GLMs (CS type × sex × genotype). Similarly, averages of context fear expression from minutes two through six of each testing day^42,63,75^ were analyzed using two-way RM GLMs within context testing sequence (test day × sex × genotype). For all data sets, *a priori* pairwise comparisons were also employed, using Bonferroni correction. One mouse (female +/−) in the cued experiment was excluded from all analyses due to failure to exceed 25% freezing for the first five CS+ tones during stage 2. One mouse (female +/+) in the context experiment was similarly excluded after failure to exceed 25% freezing during any training post-shock period. Two mice (one female +/+, one male +/−) tested for fear expression at day 28 were excluded because they failed to achieve ≥50% freezing for at least one 30 s bin. Finally, one mouse (female +/+) was excluded as an outlier, with day 28 fear expression average freezing more than 7 standard deviations below the group mean. GraphPad Prism (v 10.5.0 (673), GraphPad Software, San Diego, CA, USA) was used to generate figures showing estimated marginal means (EM means) ± 95% confidence intervals (CIs) calculated in SPSS.

## Results

None of our behavior measures produced a significant three-way interaction of time × genotype × sex. Context fear training did not reveal any interactions with genotype, but significant time × sex interactions were observed both for mice intended to be tested first in the training context followed by the novel context, and for mice undergoing the reverse testing order (Tables 1-2). No main effects of genotype were found (Tables 1-2). Mice were intentionally counterbalanced after context training to minimize the risk of acquisition confounds affecting subsequent testing responses. Pairwise comparisons supported this outcome (Fig. 1), with no genotype differences within sex at any time point, and only a single sex difference within wildtypes in the B➜A group for the first post-shock period during training (Fig 1D, E).

**Figure 1.**
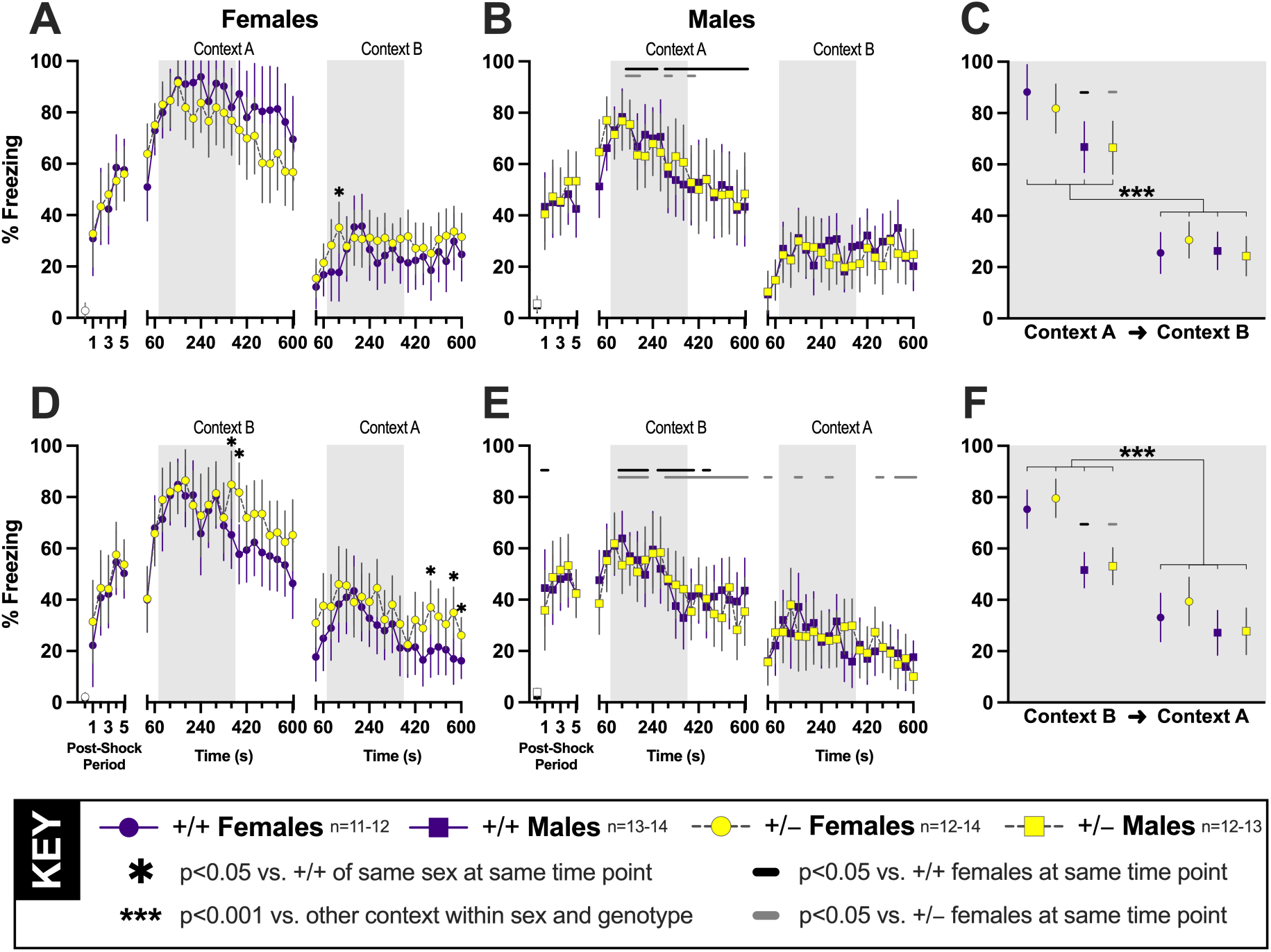
Context fear expression and generalization were tested four weeks after context fear conditioning in PMAT-deficient mice. Percent time spent freezing to the training context (Context A) or to a novel context (Context B) was measured in female (panels A, D; circles) and male (panels B, E; squares) mice. Solid purple symbols and lines indicate wildtype (+/+) data; yellow-filled grey symbols with dashed grey lines indicate heterozygous (+/−) data. All mice underwent context fear training (left third of each panel A-B, D-E), followed 28 days later (center third of each panel A-B, D-E) by testing in the training (panels A-B) or novel (panels D-E) context, then 48 h later (right third of each panel, A-B, D-E) the same mice were tested in the novel (panels A-B) or training (panels D-E) context. Averages of minutes two through six for each of the two testing days (light grey shaded rectangles, panels A-B, D-E) are presented in panels C and F by testing sequence (C: testing in A➜B; F: testing in B➜A). The first point in the first third of panels A-B, D-E represents average baseline (two min) freezing for female (circles; A, C) and male (squares; B, D) wildtype (black symbols) or heterozygous (white-filled grey symbols) prior to any foot shock administration. ✱p=0.022 (panel A), 0.038, 0.005, 0.024, 0.013, 0.044 (panel D, left to right) vs. +/+ females within the same context testing sequence and time point. Horizontal black (+/+) or grey (+/−) bars indicate significant sex differences within genotype at the respective covered time point(s). For panels A vs. B, left to right, +/+ p=0.013, 0.014, 0.024, 0.009, 0.003, 0.001, 0.005, <0.001, 0.042, 0.017, 0.007, 0.016, 0.006, 0.001, 0.025; +/− p=0.016, 0.049, 0.035, 0.043. Panel C, +/+ p=0.001, +/− p=0.034 for context A males vs. females; ***p<0.001 vs. other testing context within sex and genotype. For panels D vs. E, left to right, +/+ p=0.046, 0.036, <0.001, 0.003, 0.001, 0.024, <0.001, 0.001, <0.001, 0.041, 0.006; +/− p=<0.001, <0.001, <0.001, 0.026, <0.001, 0.007, <0.001, <0.001, 0.002, <0.001, <0.001, 0.001, 0.015, <0.001, 0.003, 0.024, 0.049, 0.011, 0.034, 0.025, 0.012, 0.001. Panel F, +/+ p<0.001 and +/− p<0.001 for context B males vs. females; ***p<0.001 vs. other testing context within sex and genotype. Data are graphed as EM means ± 95% CI.

**Table 1.** Two-way RM GLMs of context training responses for mice to be tested first in the training context (A), then in the novel context (B).

| **A→B Context Training** | **F statistic** | **p** | **$\eta_p^2$** |
| --- | --- | --- | --- |
| Time | $F_{(3,971,182.7)}=87.58$ | <0.001 | 0.656 |
| Genotype | $F_{(1,46)}=0.101$ | 0.753 | 0.002 |
| Sex | $F_{(1,46)}=0.003$ | 0.955 | 0.000 |
| Time × Genotype | $F_{(3,971,182.7)}=0.266$ | 0.898 | 0.006 |
| Time × Sex | $F_{(3,971,182.7)}=2.905$ | 0.023 | 0.059 |
| Genotype × Sex | $F_{(1,46)}=0.082$ | 0.775 | 0.002 |
| Time × Genotype × Sex | $F_{(3,971,182.7)}=0.878$ | 0.477 | 0.019 |

**Table 2.** Two-way RM GLMs of context training for mice to be tested first in the novel context (B), then in the training context (A).

| **B→A Context Training** | **F statistic** | **p** | **$\eta_p^2$** |
| --- | --- | --- | --- |
| Time | $F_{(3,269,153.6)}=87.15$ | <0.001 | 0.650 |
| Genotype | $F_{(1,47)}=0.229$ | 0.635 | 0.005 |
| Sex | $F_{(1,47)}=0.143$ | 0.707 | 0.003 |
| Time × Genotype | $F_{(3.269, 153.632)}=0.158$ | 0.936 | 0.003 |
| **Time × Sex** | **$F_{(3.269, 153.632)}=4.241$** | **0.005** | **0.083** |
| Genotype × Sex | $F_{(1,47)}=0.089$ | 0.767 | 0.002 |
| Time × Genotype × Sex | $F_{(3.269, 153.632)}=0.933$ | 0.433 | 0.019 |

As with context training, context testing responses on day 28 did not result in any genotype interactions nor a main genotype effect, but the interaction of time × sex was again significant in both testing groups (Tables 3-4). Pairwise comparisons did not indicate any significant differences across genotype in either sex for mice tested first for context fear expression, then for context fear generalization (A➜B; Fig. 1A, B). A sex difference was observed in +/+ mice, with pairwise comparisons highlighting that females exhibited greater freezing than males during 75% of day 28’s testing (Fig. 1A, B). However, PMAT deficiency attenuated most of these sex differences on day 28, with only 20% of male +/− time points being significantly less than female +/− freezing (Fig. 1A, B). In mice tested initially for context fear generalization, then for context fear expression (B➜A; Fig. 1D, E), pairwise comparisons suggest a different pattern. Both genotypes across sexes exhibited fear generalization on day 28, and reduced PMAT function transiently prolonged this generalization in females, but not males (Fig. 1D, E). This was made more apparent when comparing across sexes. While decreases in PMAT attenuated sex differences for A➜B mice, sex differences became more pronounced for B➜A +/− mice on day 28. Indeed, 75% of day 28 male +/− freezing time points were significantly less than +/− females, while only half of +/+ male timepoints were lower than +/+ females (Fig. 1D, E). Thus, extinction of fear generalization may be enhanced by intact PMAT function in females, but not in males.

**Table 3.**
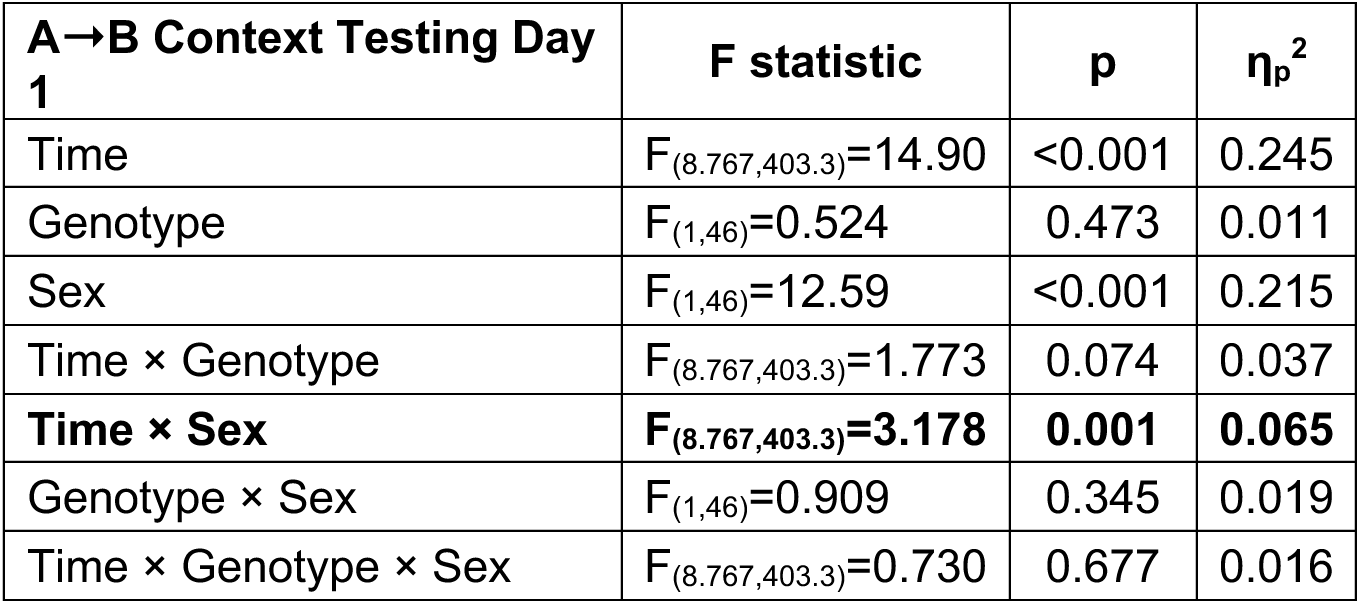
Two-way RM GLMs of context testing responses for mice tested on day 28 in the training context (A).

| **A→B Context Testing Day 1** | **F statistic** | **p** | **$\eta_p^2$** |
| --- | --- | --- | --- |
| Time | $F_{(8,767,403.3)}=14.90$ | <0.001 | 0.245 |
| Genotype | $F_{(1,46)}=0.524$ | 0.473 | 0.011 |
| Sex | $F_{(1,46)}=12.59$ | <0.001 | 0.215 |
| Time × Genotype | $F_{(8,767,403.3)}=1.773$ | 0.074 | 0.037 |
| **Time × Sex** | **$F_{(8,767,403.3)}=3.178$** | **0.001** | **0.065** |
| Genotype × Sex | $F_{(1,46)}=0.909$ | 0.345 | 0.019 |
| Time × Genotype × Sex | $F_{(8,767,403.3)}=0.730$ | 0.677 | 0.016 |

**Table 4.**
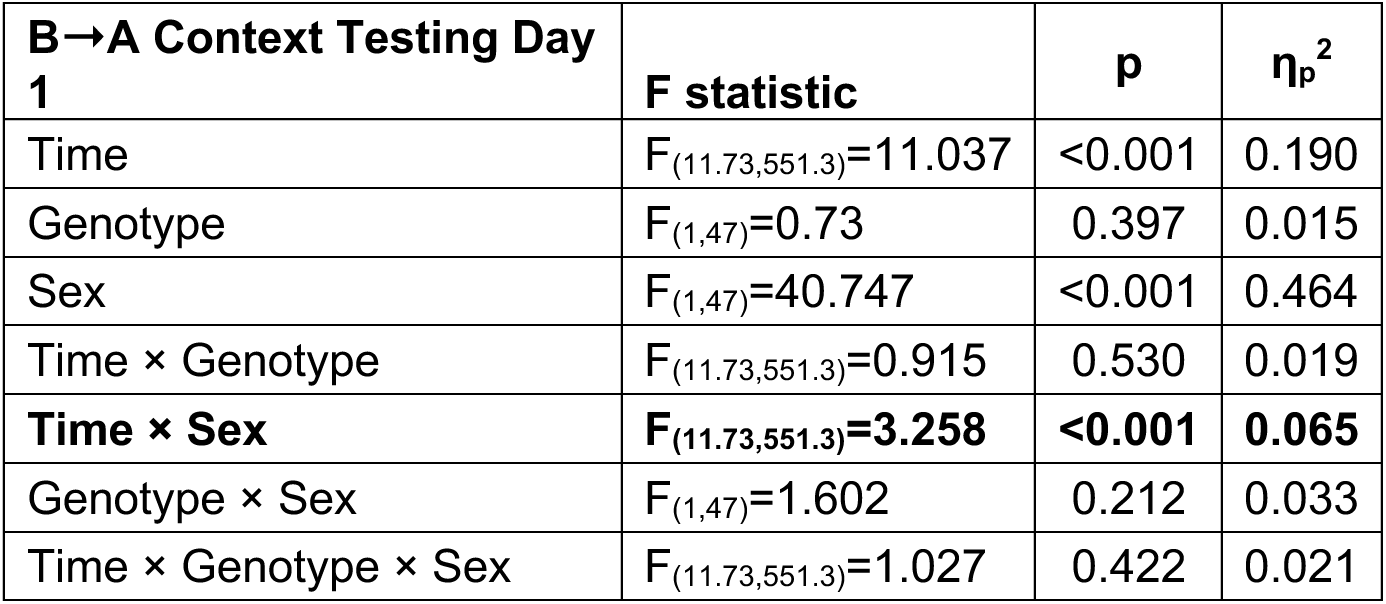
Two-way RM GLMs of context testing responses for mice tested on day 28 in the novel context (B).

Deviating slightly from the preceding patterns, day 30 context testing responses for A➜B and B➜A mice displayed no two-way interactions (Tables 5-6). Both A➜B and B➜A day 30 behavior produced significant main effects of time (Tables 5-6), and a main effect of sex was also significant for B➜A mice (Table 6). A single time point was found by pairwise comparisons to indicate elevated freezing in +/+ as compared to +/− A➜B females on day 30 (Fig. 1A), but otherwise no genotype nor sex differences were observed at any specific time points (Fig. 1A, B). Akin to day 28 testing for B➜A mice, pairwise comparisons reported sustained elevations in B➜A +/− female freezing near the end of day 30 testing (Fig. 1D), when extinction processes are typically more prominent. Moreover, female +/− mice in the B➜A testing sequence continued to exhibit elevated freezing during 35% of day 30’s time points vs. male +/− mice, whereas +/+ mice in this group displayed no significant pairwise comparisons across sex (Fig. 1D, E).

**Table 5.**
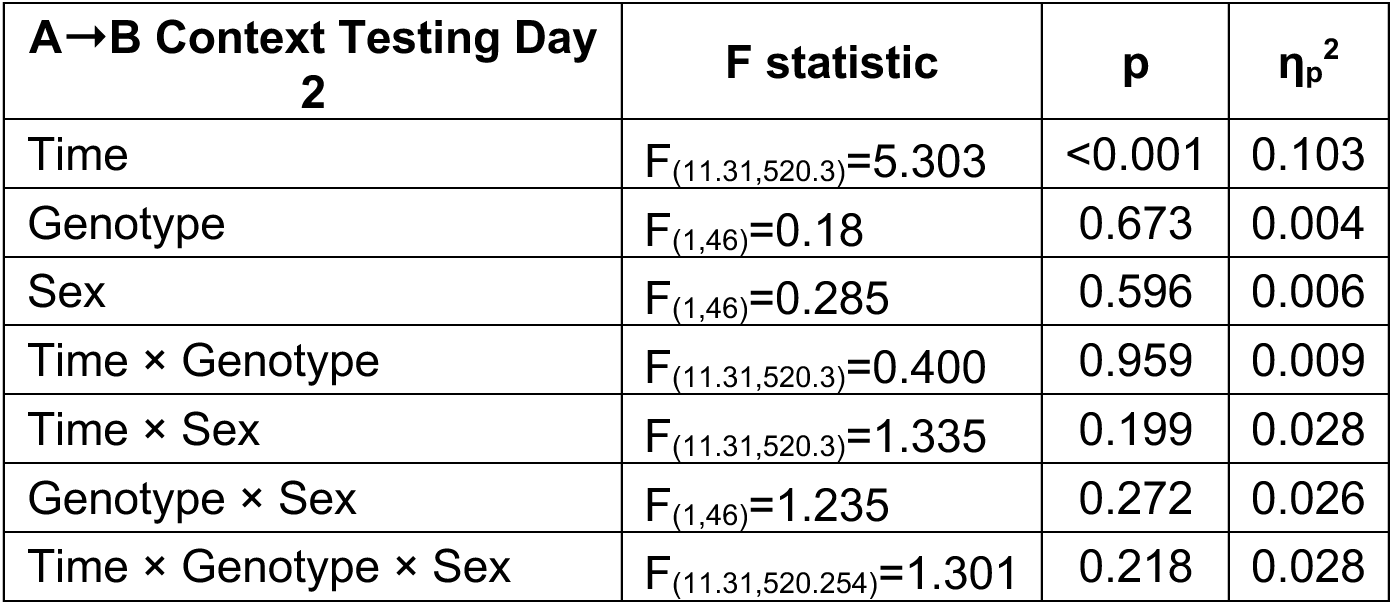
Two-way RM GLMs of context testing responses for mice tested on day 30 in the novel context (B).

**Table 6.** Two-way RM GLMs of context testing responses for mice tested on day 30 in the training context (A).

| **B→A Context Testing Day**<br>**2** | **F statistic** | **p** | **$\eta_p^2$** |
| --- | --- | --- | --- |
| Time | $F_{(9.014,423.7)}=8.007$ | <0.001 | 0.146 |
| Genotype | $F_{(1,47)}=1.636$ | 0.207 | 0.034 |
| Sex | $F_{(1,47)}=4.299$ | 0.044 | 0.084 |
| Time × Genotype | $F_{(9.014,423.7)}=1.322$ | 0.223 | 0.027 |
| Time × Sex | $F_{(9.014,423.7)}=0.949$ | 0.483 | 0.020 |
| Genotype × Sex | $F_{(1,47)}=1.346$ | 0.252 | 0.028 |
| Time × Genotype × Sex | $F_{(9.014,423.7)}=0.745$ | 0.668 | 0.016 |

To focus on initial context fear expression or generalization, while minimizing evaluation of extinction, we averaged freezing for all mice during minutes two through six on each of the two testing days^42,63,73^. Returning to the earlier noted pattern, analyses of these testing data reported significant time × sex interactions, without any genotype interactions or main effects (Tables 7-8). Pairwise comparisons upheld the sex differences within genotypes, as well as the anticipated order effect of freezing on day 30 being less than day 28 (Fig. 1C, F). A non-significant trend (p=0.080, ηp^2^=0.064) was noted between female and male +/− mice in the B➜A group on day 30 (Fig. 1F), aligning with our aforementioned observations.

**Table 7.** Two-way RM GLMs of average context testing responses during minutes two through six for mice tested first in the training context (A), then in the novel context (B).

| **A→B Context Testing Averages** | **F statistic** | **p** | **$\eta_p^2$** |
| --- | --- | --- | --- |
| Time | $F_{(1,46)}=421.9$ | <0.001 | 0.902 |
| Genotype | $F_{(1,46)}=0.061$ | 0.806 | 0.001 |
| Sex | $F_{(1,46)}=8.031$ | 0.007 | 0.149 |
| Time × Genotype | $F_{(1,46)}=1.012$ | 0.320 | 0.022 |
| **Time × Sex** | **$F_{(1,46)}=10.59$** | **0.002** | **0.187** |
| Genotype × Sex | $F_{(1,46)}=0.004$ | 0.952 | 0.000 |
| Time × Genotype × Sex | $F_{(1,46)}=1.914$ | 0.173 | 0.040 |

**Table 8.** Two-way RM GLMs of average context testing responses during minutes two through six for mice tested first in the novel context (B), then in the training context (A).

| **B→A Context Testing Averages** | **F statistic** | **p** | **$\eta_p^2$** |
| --- | --- | --- | --- |
| Time | $F_{(1,47)}=202.8$ | <0.001 | 0.812 |
| Genotype | $F_{(1,47)}=0.871$ | 0.356 | 0.018 |
| Sex | $F_{(1,47)}=25.11$ | <0.001 | 0.348 |
| Time × Genotype | $F_{(1,47)}=0.013$ | 0.910 | 0.000 |
| **Time × Sex** | **$F_{(1,47)}=12.35$** | **<0.001** | **0.208** |
| Genotype × Sex | $F_{(1,47)}=0.386$ | 0.537 | 0.008 |
| Time × Genotype × Sex | $F_{(1,47)}=0.100$ | 0.753 | 0.002 |

Distinct from context fear conditioning, where mice were counterbalanced after training across one of two testing sequences, mice that underwent cued fear conditioning here were all within the same group. Instead, all mice were presented with both CS+ and CS− auditory cues throughout training and testing; it is merely the responses to each of these that are graphed separately for clarity (Fig. 2).

**Figure 2.**
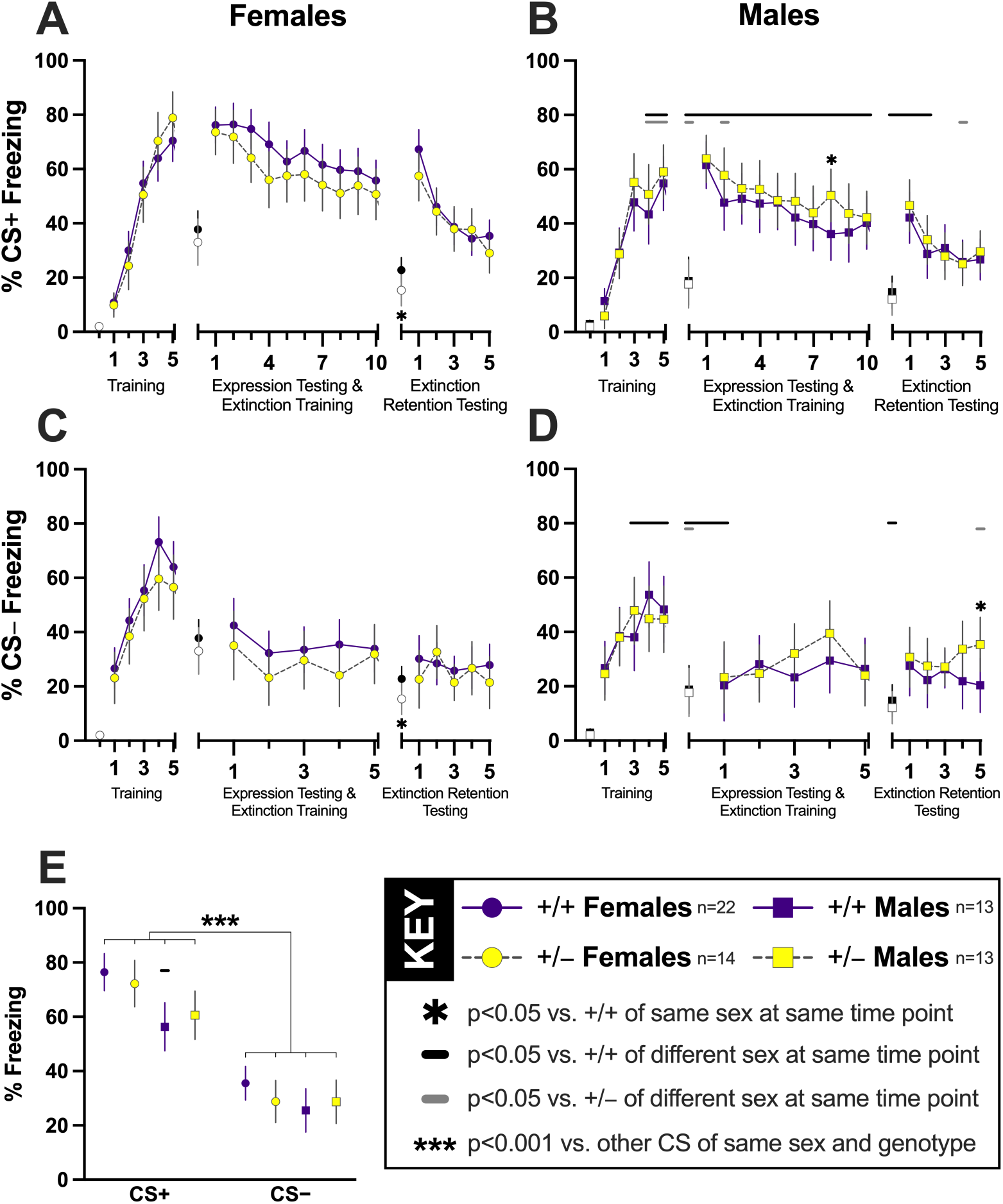
Discrimination of explicitly paired and unpaired auditory cues during fear conditioning in PMAT-deficient mice. Percent time spent freezing to the conditioned stimulus (CS) that was explicitly paired (CS+, 7.5 kHz; panels A-B) or never paired (CS−, 2.0 kHz; panels C-D) with an aversive mild foot shock are shown for female (panels A, C; circles) and male (panels B, D; squares) mice. Solid purple symbols and lines indicate wildtype (+/+) data; yellow-filled grey symbols with dashed grey lines indicate heterozygous (+/−) data. All mice underwent cued fear training (left third of each panel A-D) that employed five presentations each of the CS+ and CS−, followed 48 h later by cued expression testing and extinction training (center third of each panel A-D) involving 30 CS+ and 5 CS− tone presentations, and concluding 48 h thereafter by cued extinction retention testing (right third of each panel A-D) employing 15 CS+ and 5 CS− presentations. Freezing to CS+ on the two testing days are graphed as averages of three sequential CS+ presentations, creating 10 data points for the first testing day, and 5 for the second testing day. The first point in each third of panels A-D represents average baseline (two min) freezing for female (circles; A, C) and male (squares; B, D) wildtype (black symbols) or heterozygous (white-filled grey symbols) in the absence of any tones; thus, within sex and genotype, these baseline values are identical in panels A vs. C and B vs. D. Panel E shows the average percent time freezing during the first 5 CS+ (left half) and 5 CS− (right half) tones presented during cued expression testing and extinction training. ✱p=0.046, 0.041, 0.046 vs. +/+ of same sex within CS and time point, in alphabetical panel order. Horizontal black (+/+) or grey (+/−) bars indicate significant sex differences within genotype at the respective covered time point(s). For panels A vs. B, left to right, +/+ p=0.004, 0.015, 0.001, 0.009, <0.001, <0.001, 0.002, 0.020, <0.001, <0.001, <0.001, 0.002, 0.014, 0.034, <0.001, 0.003; +/− p=0.012, 0.006, 0.016, 0.049, 0.026. For panels C vs. D, left to right, +/+ p=0.028, 0.013, 0.047, 0.001, 0.010, 0.034; +/− p=0.016, 0.049. Panel E, +/+ p<0.001 for male vs. female CS+; ***p<0.001 vs. other CS within sex and genotype. Data are graphed as EM means ± 95% CI.

Training to the CS+, similar to many of the context fear analyses, revealed a significant interaction between time × sex (Table 9). Genotype was not a component of any significant two-way interactions nor a main effect (Table 9). As before, the time × sex interaction was expected, with females of both genotypes conditioning to the CS+ with higher freezing percentages than males, supported by pairwise comparisons (Fig. 2A, B).

**Table 9.** Two-way RM GLMs of cued training responses to the CS explicitly paired with the aversive stimulus (CS+).

| **Cued CS+ Training** | **F statistic** | **p** | **$\eta_p^2$** |
| --- | --- | --- | --- |
| **Time** | **$F_{(4,232)}=225.3$** | **<0.001** | **0.795** |
| Genotype | $F_{(1,58)}=0.322$ | 0.573 | 0.006 |
| Sex | $F_{(1,58)}=7.295$ | 0.009 | 0.112 |
| Time × Genotype | $F_{(4,232)}=1.593$ | 0.177 | 0.027 |
| **Time × Sex** | **$F_{(4,232)}=7.746$** | **<0.001** | **0.118** |
| Genotype × Sex | $F_{(1,58)}=0.079$ | 0.779 | 0.001 |
| Time × Genotype × Sex | $F_{(4,232)}=0.765$ | 0.549 | 0.013 |

Two days after training, testing for CS+ cued fear expression concurrent with CS+ extinction training resulted in no two-way interactions, but significant main effects of both sex and time (Table 10). These again support the anticipated greater freezing levels in females vs. males, along with time-dependent reductions in freezing across sexes indicative of extinction (Table 10; Fig. 2A, B). Pairwise comparisons of each genotype within sex indicated a single time point where +/− males displayed more freezing than +/+ males (Fig. 2B). More informative are the pairwise comparisons across sexes at each genotype’s timepoint. Similar to context A➜B day 28 testing, +/+ males on the first day of cued testing displayed less freezing than +/+ females at all analyzed CS+ measures, whereas +/− males exhibited lower CS+ fear compared to +/− females for a mere 18% of that day’s measures (Fig. 2A, B). In other words, PMAT function may enhance sex differences in cued fear discrimination of the CS+.

**Table 10.**
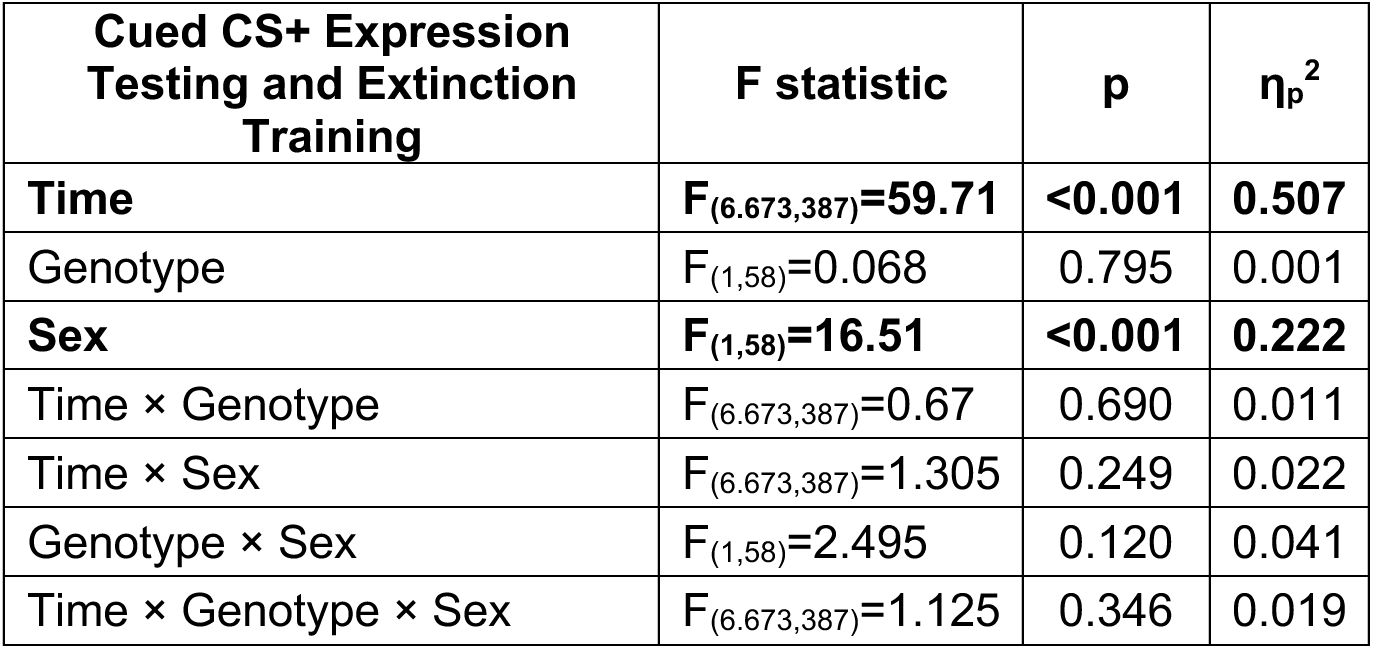
Two-way RM GLMs of cued CS+ expression testing and extinction training responses.

As with cued CS+ training, a significant interaction of time × sex was found without any genotype interactions nor a main genotype effect on the final cued testing day evaluating retention of cued fear extinction training (Table 11). Pairwise comparisons indicated that +/− females had less baseline freezing at the start of this extinction retention test than +/+ females (Fig. 2A, C), the sole instance in this study where PMAT deficiency statistically attenuated fear behavior. The pattern of sex differences being more prominent between +/+ as opposed to +/− mice only partially (50%) persisted on this last cued testing day, disappearing for the last three CS+ measures for +/+ mice (Fig. 2A, B). A single CS+ measure in +/− mice occurred where females froze more than males (Fig. 2A, B).

**Table 11.** Two-way RM GLMs of cued CS+ extinction retention testing responses.

| **Cued CS+ Extinction Retention Testing** | **F statistic** | **p** | **$\eta_p^2$** |
| --- | --- | --- | --- |
| **Time** | **$F_{(4.347,252.1)}=107.5$** | **<0.001** | **0.650** |
| Genotype | $F_{(1,58)}=0.189$ | 0.666 | 0.003 |
| Sex | $F_{(1,58)}=10.17$ | 0.002 | 0.149 |
| Time × Genotype | $F_{(4.347,252.1)}=1.179$ | 0.321 | 0.020 |
| **Time × Sex** | **$F_{(4.347,252.1)}=4.900$** | **<0.001** | **0.078** |
| Genotype × Sex | $F_{(1,58)}=0.574$ | 0.452 | 0.010 |
| Time × Genotype × Sex | $F_{(4.347,252.1)}=2.230$ | 0.061 | 0.037 |

Additional evidence that PMAT function augments biological sex differences in fear behaviors is observed in freezing behaviors during CS− presentations. For cued CS− ‘training’, though no two-way interactions with, nor main effect of, genotype were observed, the interaction of time × sex was once more significant (Table 12). Pairwise comparisons found that for the last three measures of freezing to the CS−, +/+ but not +/− mice displayed sex differences, with males freezing less than females (Fig. 2C, D).

**Table 12.** Two-way RM GLMs of cued ‘training’ responses to the CS never paired with the aversive stimulus (CS−).

| **Cued CS- Training** | **F statistic** | **p** | **$\eta_p^2$** |
| --- | --- | --- | --- |
| **Time** | **$F_{(4,071,236.1)}=87.38$** | **<0.001** | **0.601** |
| Genotype | $F_{(1,58)}=1.493$ | 0.227 | 0.025 |
| Sex | $F_{(1,58)}=7.271$ | 0.009 | 0.111 |
| Time × Genotype | $F_{(4,071,236.1)}=1.211$ | 0.307 | 0.020 |
| **Time × Sex** | **$F_{(4,071,236.1)}=2.966$** | **0.020** | **0.049** |
| Genotype × Sex | $F_{(1,58)}=0.709$ | 0.403 | 0.012 |
| Time × Genotype × Sex | $F_{(4,071,236.146)}=0.280$ | 0.894 | 0.005 |

The absence of a sex difference in +/−, but not +/+, mice persisted through the first CS− presentation on the cued expression testing and extinction training day as revealed by pairwise comparisons (Fig. 2C, D). Also as with the CS− ‘training’ day, the only significant two-way interaction was that of time × sex, and the main effect of genotype was not significant (Table 13).

**Table 13.** Two-way RM GLMs of cued CS− expression testing and extinction training responses.

| **Cued CS- Expression Testing and Extinction Training** | **F statistic** | **p** | **$\eta_p^2$** |
| --- | --- | --- | --- |
| Time | $F_{(4,591,266.3)}=0.872$ | 0.493 | 0.015 |
| Genotype | $F_{(1,58)}=0.316$ | 0.576 | 0.005 |
| Sex | $F_{(1,58)}=4.103$ | 0.047 | 0.066 |
| Time × Genotype | $F_{(4,591,266.3)}=0.437$ | 0.807 | 0.007 |
| **Time × Sex** | **$F_{(4,591,266.3)}=4.111$** | **0.002** | **0.066** |
| Genotype × Sex | $F_{(1,58)}=1.623$ | 0.208 | 0.027 |
| Time × Genotype × Sex | $F_{(4,591,266.3)}=0.794$ | 0.545 | 0.014 |

When CS− extinction retention testing was evaluated, no two-way interactions emerged as significant. The only main effect was that of time (Table 14). Pairwise comparisons indicated an unanticipated elevation in freezing during the final CS− presentation in +/− males as compared to +/+ males (Fig. 2D), possibly indicating an emerging sensitization to the CS−. This time point is also the single instance here where male +/− mice exhibited heightened freezing behavior vs. +/− females.

**Table 14.**
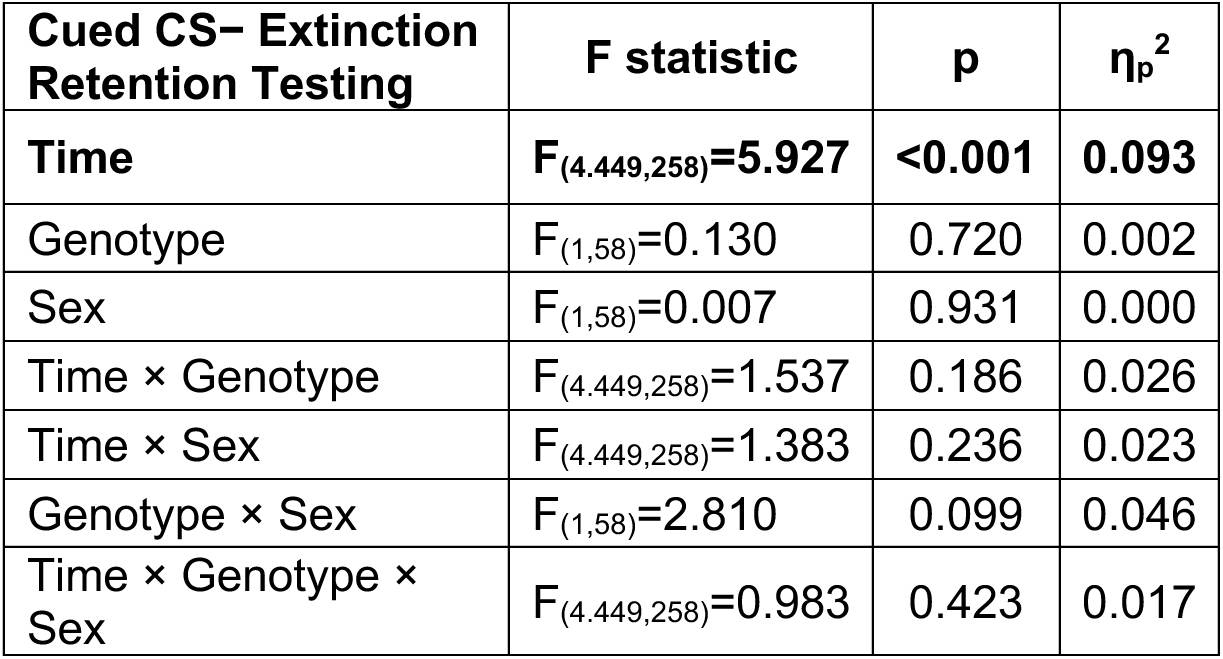
Two-way RM GLMs of cued CS− extinction retention testing responses.

| **Cued CS– Extinction Retention Testing** | **F statistic** | **p** | **$\eta_p^2$** |
| --- | --- | --- | --- |
| **Time** | **$F_{(4,449,258)}=5.927$** | **<0.001** | **0.093** |
| Genotype | $F_{(1,58)}=0.130$ | 0.720 | 0.002 |
| Sex | $F_{(1,58)}=0.007$ | 0.931 | 0.000 |
| Time × Genotype | $F_{(4,449,258)}=1.537$ | 0.186 | 0.026 |
| Time × Sex | $F_{(4,449,258)}=1.383$ | 0.236 | 0.023 |
| Genotype × Sex | $F_{(1,58)}=2.810$ | 0.099 | 0.046 |
| Time × Genotype × Sex | $F_{(4,449,258)}=0.983$ | 0.423 | 0.017 |

Related to our focus on minutes two through six of context fear testing to more selectively assess fear expression while minimizing extinction processes, for cued fear we analyzed the average of the first five CS+ and five CS− tones played during day 28’s testing of cued fear expression^66^. This facilitated a clearer analysis of cued fear discrimination versus generalization while still permitting sex and genotype analyses. Mirroring the themes noted previously, time × sex was the only significant interaction, and no main effect of genotype was found (Table 15).

**Table 15.** Two-way RM GLMs of the average responses to the first five CS+ and five CS− tones played during cued expression testing and extinction training.

| **First 5 CS+ and CS-**<br>**Cued Fear Expression** | **F statistic** | **p** | **$\eta_p^2$** |
| --- | --- | --- | --- |
| **Time** | **$F_{(1,58)}=219.9$** | **<0.001** | **0.791** |
| Genotype | $F_{(1,58)}=0.078$ | 0.782 | 0.001 |
| Sex | $F_{(1,58)}=11.42$ | 0.001 | 0.164 |
| Time × Genotype | $F_{(1,58)}=0.138$ | 0.712 | 0.002 |
| **Time × Sex** | **$F_{(1,58)}=4.784$** | **0.033** | **0.076** |
| Genotype × Sex | $F_{(1,58)}=2.212$ | 0.142 | 0.037 |
| Time × Genotype × Sex | $F_{(1,58)}=0.020$ | 0.887 | 0.000 |

Pairwise comparisons likewise illustrated clear sex differences in CS+ freezing responses across +/+ mice (Fig. 2E), and further indicated a non-significant trend (p=0.053, ηp^2^=0.063) across sexes for CS− responses in +/+ mice. Nevertheless, regardless of sex or genotype combination, all mice significantly discriminated between the CS+ and CS− tones. Thus, intact PMAT function does not appear to contribute to fear discrimination nor generalization processes as we hypothesized, but instead to canonical sexual dimorphism in freezing responses across context and cued fear conditioning paradigms.

## Discussion

Given our previous evaluations of how constitutively reduced PMAT function affects behavioral responses to environmental stressors^41,43^ including fear conditioning^42^, we anticipated sex-selective effects of PMAT deficiency here. PMAT deficiency elevates females’ active coping^43^ while suppressing the same in males^41^ during distinct inescapable stressors. Testing 48 h after training indicated that PMAT likely attenuates males’ context fear and enhances extinction of females’ cued fear following conditioning to a single auditory cue^42^. Collectively, these outcomes led to us hypothesizing that +/− males would exhibit enhanced context fear generalization, whereas +/− females would display augmented cued fear generalization. Our data instead demonstrated that PMAT appears to function as a partial suppressor of context fear generalization extinction and subsequent specificity extinction, while also enhancing behavioral differences in cued and context fear responses across biological sexes.

Prior work found +/− males exhibited sustained context fear expression compared to +/+ males when tested two days after training, indicating PMAT typically helps suppress males’ recently learned context fear behaviors^42^. Loss of PMAT function also elevates males’ passive coping behaviors^41^, providing further evidence that uptake by PMAT moderates males’ stress responses. Here, we intentionally tested context fear behaviors 4 weeks after training to evaluate context fear generalization^63,74,75^. Rather than finding our hypothesized enhancement of +/− males’ context fear generalization, we instead discovered evidence indicating PMAT contributes to extinction of context fear generalization selectively in +/− females. Moreover, +/− females’ experience of this generalization appeared to influence subsequent testing in the original training context by further impeding context-specific fear extinction. Critically, these effects were not observed in males undergoing the same testing sequence, nor were they found when mice instead were first tested in the original training context prior to generalization testing. Combined, our data suggest reduced PMAT function in biological females impedes generalization extinction and disrupts ensuing extinction processes when the initial, specific fear memory is retrieved. Whether this is through shared or distinct mechanisms requires additional investigation. Regardless, this aligns with our previous report implicating PMAT as a contributor to females’ cued fear extinction following training to a sole, 4 kHz tone with a larger magnitude aversive stimulus (0.8 mA^42^).

To further investigate generalization and discrimination in PMAT deficient mice here, we used a two tone (2 and 7.5 kHz) cued fear conditioning paradigm^66^ with a lower magnitude aversive stimulus (0.4 mA). Because PMAT appears to dampen females’ behavioral stress responses^43^ and facilitate their cued fear extinction^42^, we expected +/− females to exhibit more cued fear generalization (i.e., less cued fear discrimination) than +/− males and wildtypes across sexes. Instead, PMAT reductions did not seem to directly influence discrimination between CS+ and CS− tones, but they did consistently mitigate the sex differences observed between +/+ and +/− mice. This was not selective to auditory tone responses either, as a similar pattern was discovered when testing A➜B mice in the training context. Thus, PMAT’s augmentation of these sexual dimorphisms appears to occur when first the specific context or cue is presented, followed thereafter by the similar context or cue that can elicit generalization. Detection of this pattern across fear conditioning paradigms here, but not in our previous report, is expected given the numerous intentional methodological differences between the two studies. These include the intentional 4 week testing delay (context fear generalization) and two tone conditioning (cued fear discrimination/generalization), plus the lower foot shock intensities (0.4 vs. 0.8 mA) and inclusion of sex here as a factor in omnibus statistics (analyses were within sex for the sake of another experimental factor in^42^). While our observations of PMAT’s effects here might initially appear modest, the overarching consistency within and across studies means PMAT function could influence anxiety disorder risk. This appears particularly relevant to people assigned female at birth who have at least one of the *SLC29A4* polymorphisms associated with decreased PMAT function^37,38,40,47,48^. Identification of the sex hormone(s) contributing to these sex-specific effects, combined with determination of the organizational vs. activational origin of PMAT’s influences, could provide a substantial advancement in understanding PMAT function.

Further complicating PMAT’s sex-selective conditioned fear effects are studies reporting that interrupting dopamine or serotonin signaling attenuates fear conditioning^76,77^, sometimes in only one sex^78^. Dopamine receptor agonism similarly affects female-specific extinction retention, directionally dependent upon circulating endogenous sex hormone levels^79^. Sex-inclusive studies of dopamine and serotonin signaling contributions to fear conditioning remain relatively sparse. Still, the established contributions of these monoamines to high arousal situations^1–6^, combined with PMAT’s preferential and accelerated transport of dopamine and serotonin^21,23,27^, make it probable that PMAT contributes to additional, translationally meaningful, neurophysiological and behavioral stress responses.

Limitations of our study include the use of mice constitutively deficient in PMAT, meaning lifelong compensatory processes may have been engaged. However, we intentionally used only +/+ and +/−, and not −/−, mice, to both minimize this potential confound and to maximize the translatability of our findings to humans with one or more of the many *SLC29A4* functional polymorphisms^37,38,40,47,48^. The precise magnitude of functional PMAT reduction in +/− mice remains to be quantified, in part because spatial quantification methods like autoradiography are precluded due to the absence of drugs selective for PMAT^28–30^. Commercial PMAT antibody reliability remains too inconsistent at present for a diligent assessment of brain-wide +/− reductions in PMAT levels^80^, particularly considering the spatial nuances of PMAT expression^81^. Moreover, measurement of PMAT expression in mice +/− mice would not necessarily translate to functional decreases, returning again to the limitation of absent pharmacological tools. We intentionally did not monitor estrous cycles in our female mice, because we wanted to observe outcomes in naturally cycling females, and because multiple labs have demonstrated that C57BL/6J mice’s estrous cycles do not affect fear conditioning^65,82,83^.

Our findings partially supported our hypotheses, plus provided additional unanticipated insights into probable contributions of PMAT function upon translationally relevant processes regulating learned aversive associations. Given the current absence of selective pharmacological tools that could advance study of PMAT^28–30^, constitutive PMAT deficient mice remain the best way to study PMAT function – via subtraction – on an organismal level. Growing evidence from human *SLC29A4* polymorphism association studies^30,37,38,40,47,48^ underscore how functional PMAT reductions exert effects upon an underappreciated breadth of physiological and behavioral outcomes. Here, our murine data indicate intact PMAT function promotes females’, but not males’, extinction of both generalized and subsequent context-specific fear. Further, functional PMAT may contribute to sex differences in long-term (i.e., remote) context fear expression^63,65,83–85^) and to auditory cues encountered during fear conditioning (whether explicitly or never paired with aversive stimuli)^66,86–89^. In people, this means genetically attenuated PMAT function might put those assigned female at birth at greater risk for anxiety disorders involving sustained, pathological fear generalization over months or years^52,90^. Future evaluations of how functional PMAT polymorphisms are associated with psychological measures of generalized anxiety, perceived stress, emotion processing, and other high arousal dimensions would be informative. Such investigations might even lead to inclusion of *SLC29A4* polymorphisms in pharmacogenomic evaluations for neuropsychiatric disorder risk and/or psychoactive drug treatment responses.

## Acknowledgements

This work was supported by Kent State University, including summer undergraduate research experience internships and Brain Health Research Institute (BHRI) summer undergraduate fellowships through the Office of Student Research and BHRI, respectively, for both AJR and LRS. This project, and additional support for LRS, were also provided by a BHRI Gold Award and University Research Council supplemental funds to LMG. We are grateful to our expert veterinarians and vivarium staff, and respectfully acknowledge the lives of the mice used in our research.

## Notes

### Competing Interest Statement

The authors have declared no competing interest.

